# Single-cell Transcriptome of Bronchoalveolar Lavage Fluid Reveals Dynamic Change of Macrophages During SARS-CoV-2 Infection in Ferrets

**DOI:** 10.1101/2020.11.18.388280

**Authors:** Jeong Seok Lee, June-Young Koh, Kijong Yi, Young-Il Kim, Su-Jin Park, Eun-Ha Kim, Se-Mi Kim, Sung Ho Park, Young Seok Ju, Young Ki Choi, Su-Hyung Park

## Abstract

Although the profile of immune cells changes during the natural course of SARS-CoV-2 inflection in human patients, few studies have used a longitudinal approach to reveal their dynamic features. Here, we performed single-cell RNA sequencing of bronchoalveolar lavage fluid cells longitudinally obtained from SARS-CoV-2-infected ferrets. Landscape analysis of the lung immune microenvironment showed dynamic changes in cell proportions and characteristics in uninfected control, at 2 days post-infection (dpi) (early stage of SARS-CoV-2 infection with peak viral titer), and 5 dpi (resolution phase). NK cells and CD8^+^ T cells exhibited activated subclusters with interferon-stimulated features, which were peaked at 2 dpi. Intriguingly, macrophages were classified into 10 distinct subpopulations, and their relative proportions changed over the time. We observed prominent transcriptome changes among monocyte-derived infiltrating macrophages and differentiated M1/M2 macrophages, especially at 2 dpi. Moreover, trajectory analysis revealed gene expression changes from monocyte-derived infiltrating macrophages toward M1 or M2 macrophages and identified the distinct macrophage subpopulation that had rapidly undergone SARS-CoV-2-mediated activation of inflammatory responses. Finally, we found that different spectrums of M1 or M2 macrophages showed distinct patterns of gene modules downregulated by immune-modulatory drugs. Overall, these results elucidate fundamental aspects of the immune response dynamics provoked by SARS-CoV-2 infection.

## Introduction

During the current coronavirus disease-19 (COVID-19) pandemic ^1^, cross-sectional research has rapidly broadened our understanding of the immune response to severe acute respiratory syndrome coronavirus 2 (SARS-CoV-2). Immune landscape studies have revealed the pathogenesis of severe COVID-19 with a hyper-inflammatory response^2–4^, and the innate, humoral, and T-cell response of COVID-19 patients have been extensively characterized^5–7^. Currently ongoing studies are examining the mechanisms of therapeutic modalities, including anti-viral and anti-inflammatory agents, with accompanying clinical trials ^8–11^. However, due to the intrinsic limitations of observational studies of human subjects, it is rare to obtain a longitudinal description of the immune response from the initial stage to the resolution of SARS-CoV-2 infection.

In recent studies, single-cell RNA sequencing (scRNA-seq) of bronchoalveolar lavage (BAL) fluid from patients with COVID-19 has provided valuable information of the microenvironment of immune responses to SARS-CoV-2 ^12-14^. Intriguingly, increased levels of a macrophage subtype originated from circulating monocytes were observed during the inflammatory phase of COVID-19 ^12^. Additionally, we recently demonstrated that peripheral monocytes from severe COVID-19 patients were highly activated, showing strong interferon-mediated inflammatory responses ^4^. These findings suggest that both monocytes and macrophages are major cell population of interest in COVID-19 pathogenesis and patients’ anti-viral response. However, most currently available transcriptomic analyses of immune cells are from cross-sectional studies and, importantly, cannot compare infected status with uninfected status due to the lack of data obtained prior to the SARS-CoV-2 infection. Moreover, BAL invasiveness hinders the acquisition of sequential specimens from critical patients during SARS-CoV-2 infection. These limitations can be overcome by analyzing animal models for the infection with SARS-CoV-2.

The ferret (*Mustela putorius furo*) is widely used as an animal model for investigations of respiratory virus pathogenesis 15,16. Since ferrets’ natural susceptibility to influenza virus was discovered in 1933, these animals have been used to recapitulate the course of several human respiratory viral diseases, including parainfluenza virus, respiratory syncytial virus, and SARS-CoV-1^17^. Moreover, their histoanatomical features—including the ratio between the upper and lower respiratory tract lengths, airway glandular density and terminal bronchiole structure—provide optimal conditions for mimicking human respiratory infection ^17^. We recently reported that a ferret model can reproduce a common natural course of COVID-19 in humans, showing effective infection and rapid transmission^18^. SARS-CoV-2-infected ferrets initially exhibit body temperature elevation and weight loss with viral shedding. In addition, peak viral titer is observed during 2-4 days post-infection (dpi), and after then, resolution phase which is characterized by body temperature normalization and decrease of viral titer is continued up to 10 days.

Here, we performed scRNA-seq of sequential BAL fluid samples from SARS-CoV-2-infected ferrets, in negative control, at 2 days post-infection (dpi) (early stage of SARS-CoV-2 infection with peak viral titer), and 5 dpi (resolution phase with histopathology). Landscape analysis of the ferret lung immune microenvironment revealed dynamic changes of the proportions and characteristics of immune cells over this time. Specifically, we delineated the macrophage population into 10 distinct subpopulations based on unique gene expression patterns, and described their chronological transcriptome changes. Intriguingly, rather than tissue-resident alveolar macrophage populations, we found that infiltrating macrophages could differentiate into M1 or M2 macrophages after SARS-CoV-2 infection. Moreover, the different spectrums of M1 or M2 macrophages exhibited distinct patterns of gene modules down-regulated by immune-modulatory drugs.

## Results

### Single-Cell Transcriptomes of BAL Fluid Cells Sequentially Obtained From SARS-CoV-2-Infected Ferrets

Ferrets were intranasally inoculated with SARS-CoV-2, using a previously described strain isolated from a COVID-19 patient in South Korea^18^. BAL fluid cells and contralateral lung tissue samples were collected by sacrificing infected ferrets at three different time-points: before SARS-CoV-2 infection (uninfected control, n = 3), 2 dpi (n = 3), and 5 dpi (n = 4) (Fig. 1a).

**Fig. 1.**
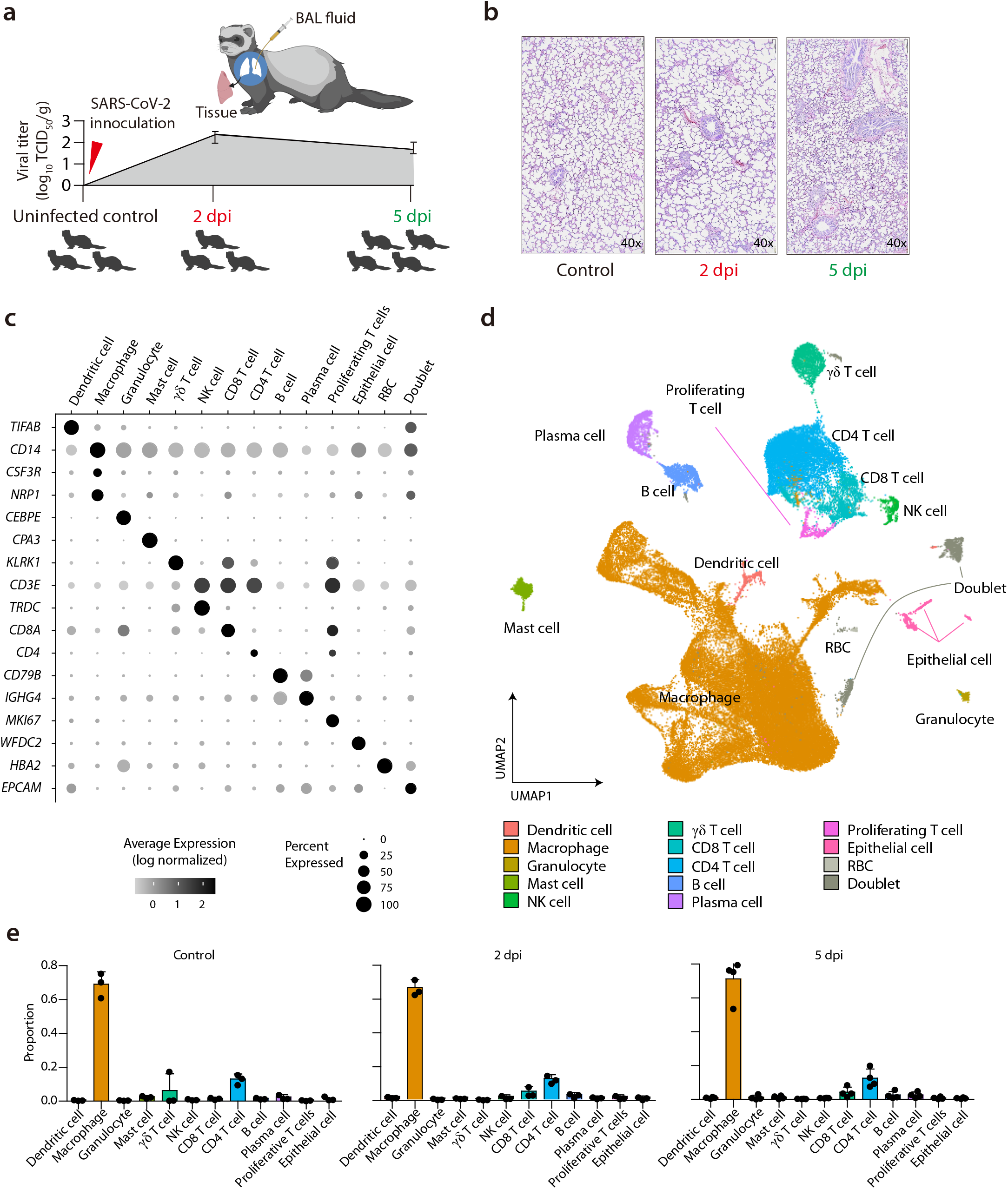
Single-Cell Transcriptomes of Bronchoalveolar Lavage (BAL) Fluid Cells From SARS-CoV-2-Infected Ferrets. a. Summary of experimental conditions with viral titers in negative control, at 2 days post-infection (dpi) and 5 dpi. b. Lung tissues of ferrets in negative control, at 2 dpi and 5 dpi with SARS-CoV-2. c. Fourteen different clusters and their specific marker gene expression levels, where brightness indicates log-normalized average expression, and circle size indicates percent expressed. d. UMAP of 59,138 cells from the BAL fluid of 10 ferrets, colored to show annotated cell types. e. Proportion of each cell type at 0 dpi (n = 3), 2 dpi (n = 3), and 5 dpi (n = 4).

Histopathological analysis and viral shedding clearly indicated SARS-CoV-2 infection (Fig. 1a and 1b). The infectious viruses detected in lung tissue at 2 dpi (mean 2.3 log_10_ TCID_50_/g) and 5 dpi (mean 1.6 log_10_ TCID_50_/g). Histopathological examinations revealed a pattern of acute pneumonia, characterized by more prominent immune cell infiltration in the alveolar wall and bronchial epithelium at 5 dpi than control or 2 dpi, which is consistent with our recent study^18^. Therefore, we categorized the 2 dpi specimens as early stage of SARS-CoV-2 infection with peak viral titer, while 5 dpi specimens may represent as resolution phase with decreasing viral titer and evident histopathological changes.

Using the 10x Genomics platform, we performed scRNA-seq of BAL fluid cells from 10 ferrets, analyzing a total of 59,138 cells after filtering dead cells. We detected a mean of 8,760 UMIs, and an average of 2,158 genes per cell. By analyzing 59,138 cells with a uniform manifold approximation and projection (UMAP) algorithm based on variable genes with the Seurat package^19^, we identified 28 different clusters (Supplementary Fig. 1a), which were assigned to 14 different cell types expressing representative marker genes (Fig. 1c, Supplementary Fig. 1b and 1c; Supplementary Table 1). We excluded two clusters with doublet and red blood cells, and thus focused on the following 12 clusters for downstream analysis: dendritic cells, macrophages, granulocytes, mast cells, natural killer (NK) cells, γδ-T cells, CD8^+^ T cells, CD4^+^ T cells, proliferating T cells, B cells, plasma cells, and epithelial cells (Fig. 1d). These clusters and annotated cell types were unbiased according to experimental batches of scRNA-seq (Supplementary Fig. 1d). Although the SARS-CoV-2 RNA sequence was rarely detected, they were contained by the macrophage and epithelial cell clusters (Supplementary Fig. 1e).

To analyze the time-course and dynamic changes of immune responses to SARS-CoV-2, we compared the relative proportions of each cell type in control, 2 dpi, and 5 dpi. Analyzing the pattern of proportion changes revealed that the macrophage population comprised the majority of BAL fluid cells over 60% (Fig. 1e). Pattern of each cell type proportion was not evidently changed regardless of time point (Fig. 1e).

### Quantitative and Qualitative Changes in the Clusters of NK Cells and CD8^+^ T Cells

As we aimed to investigate immunological changes during the early stage of SARS-CoV-2 infection, we first analyzed NK cells, the representative innate cytotoxic lymphocytes in anti-viral response. Among NK cells, five subclusters were identified from UMAP (Fig. 2a; Supplementary table 2). With regards to the proportions of each NK cluster, NK cluster 0 was decreased after SARS-CoV-2 infection, NK cluster 1 was increased at 2 dpi but decreased at 5 dpi, and NK clusters 2 and 3 were reciprocally changed (Fig. 2b and Supplementary Fig. 2a). To characterize activated status of each NK cluster, we performed gene set enrichment analysis using interferon (IFN)-α or IFN-γ responsive signatures. NK clusters 0 and 1 featured prominent responses to interferon IFN-α or IFN-γ (Supplementary Fig. 2b). Indeed, NK cluster 1 showed predominant expression of IFN-stimulated genes including *STAT1*, *OAS1*, and *ISG15* (Fig. 2c). In addition, genes of cytotoxic molecules including *GZMB*, *GZMK*, and *PRF1* were also highly expressed (Fig. 2c)—indicating that NK cluster 1 was IFN-stimulated and activated NK cells. Collectively, NK cell cluster exhibited activated subclusters with IFN-stimulated and cytotoxic features, which underwent longitudinal changes peaked at 2 dpi.

**Fig. 2.**
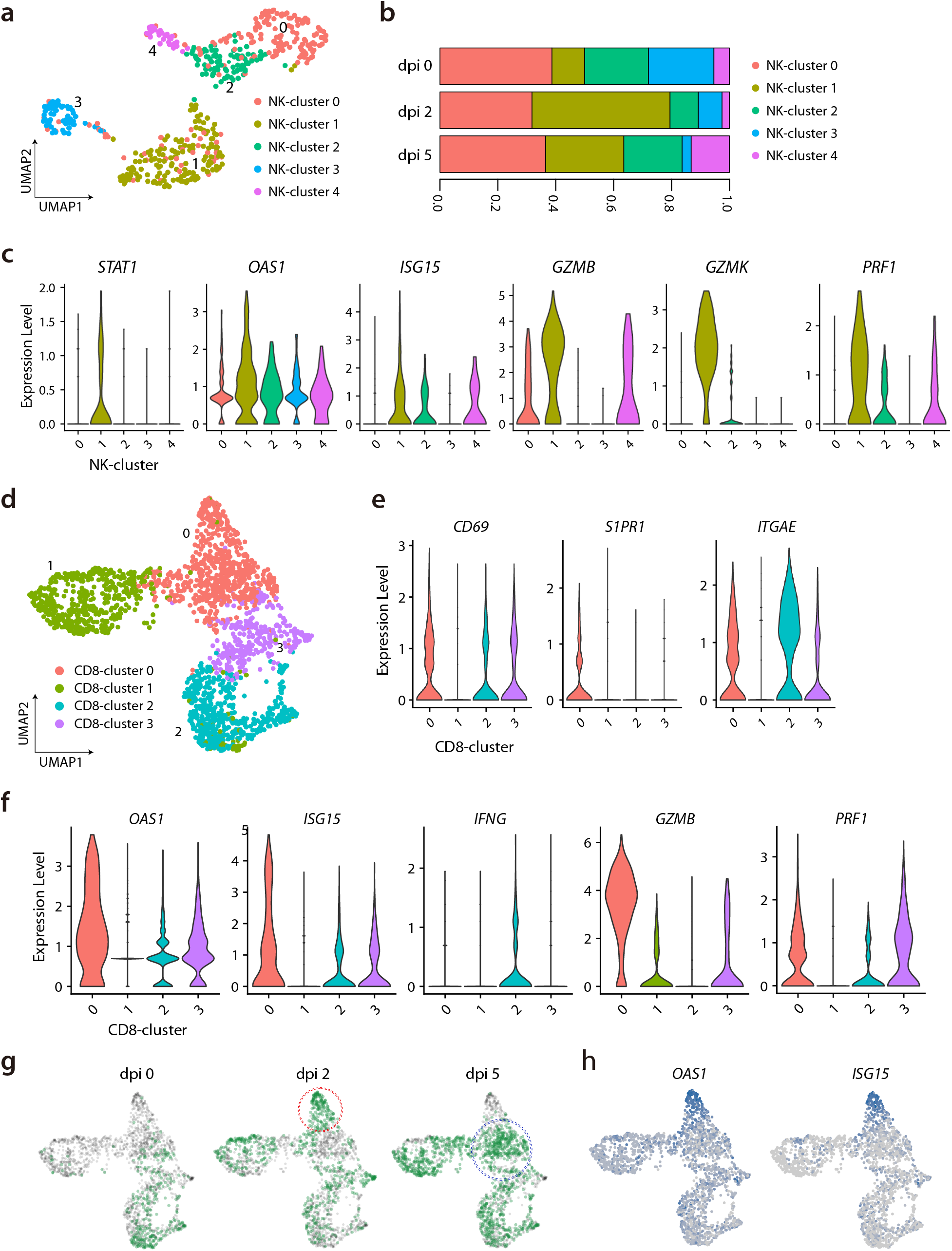
Subpopulation Analysis of NK Cells and CD8+ T Cells. a. UMAP plot of the NK cell subpopulations in all groups, colored to indicate cluster information. b. Proportion of each cell type in NK cell clusters at 0 dpi (n = 3), 2 dpi (n = 3), and 5 dpi (n = 4). c. Violin plots showing expression levels of STAT1, OAS1, ISG15, GZMB, GZMK, and PRF1 in the five NK cell clusters. d. UMAP plot of the CD8+ T-cell subpopulations in all groups, colored to show cluster information. e, f. Violin plots showing expression levels of CD69, S1PR1, ITGAE, OAS1, ISG15, IFNG, GZMB, and PRF1 in the four CD8+ T cell clusters. g. UMAP plot in which color density reflects the distributions of CD8+ T cells ferrets in negative control, at 2 dpi and 5 dpi with SARS-CoV-2. Red circle indicates concentrated area of cluster 0 with CD8^+^ T cells at 2 dpi, and blue circle indicates that of CD8^+^ T cells at 5 dpi. h. UMAP plots show normalized expressions of OAS1 and ISG15 in CD8^+^ T cells.

Additionally, we analyzed CD8^+^ T cells, another cytotoxic lymphocyte population, and identified four subclusters from UMAP (Fig. 2d; Supplementary table 3). The proportion of CD8^+^ cluster 2 tended to decrease at 2 dpi and to increase at 5 dpi, while the proportion of CD8^+^ cluster 0 reciprocally changed (Supplementary Fig. 2c). When we characterize each CD8^+^ cluster, CD8^+^ clusters 2 and 3 exhibited higher expression levels of *CD69* and *ITGAE*, and lower expression of *S1PR1*, reflecting tissue-resident features (Fig. 2e). CD8^+^ cluster 2 showed higher expressions of *CD69* and *ITGAE*, as well as high expression of *IFNG*. These findings were consistent with human CD8^+^ resident memory T (T_RM_) cells, which rapidly induce quick IFN-γ production using preformed mRNA^20^. Similar to NK cluster 1, CD8^+^ cluster 0 exhibited prominent expression of IFN-stimulated genes (including *OAS1* and *ISG15*) and the genes of cytotoxic molecules (including *GZMB* and *PRF1*) (Fig. 2f). These findings indicated that CD8^+^ cluster 0 comprised activated CD8^+^ cells; however, these cells expressed scarce amounts of *IFNG*. CD8^+^ cluster 0 showed different distributions at 2 dpi (red circle) and 5 dpi (blue circle) (Fig. 2g), which was reflected by higher IFN-stimulated signatures, including *OAS1* and *ISG15* at 2 dpi (Fig. 2h).

### Macrophage Populations Underwent Dynamic Changes According to the Natural Course of SARS-CoV-2 Infection

We next studied macrophage-specific features that dynamically changed during SARS-CoV-2 infection, since macrophage was consistently comprised the majority of cell proportion regardless of time point (Fig. 1e). To this end, we performed sub-clustering analysis of the macrophage cluster depicted in Fig. 1d. To annotate cell types, we analyzed 40,241 cells using the UMAP algorithm based on variable genes with the Seurat package^19^, and identified 17 different sub-clusters (Supplementary Fig. 3a). Based on signature genes, we selected the following 10 macrophage clusters for downstream analysis: resting tissue macrophages, APOE^+^ tissue macrophages, activated tissue macrophages, SPP1^hi^CHIT1^int^ profibrogenic M2, monocyte-derived infiltrating macrophages, weakly activated M1 macrophages, highly activated M1 macrophages, proliferating macrophages, engulfing macrophages, and unclassified cells (Fig. 3a, 3b, and Supplementary Fig. 3b). Table S4 lists the specific markers used to define each macrophage sub-cluster. Supplementary Fig. 3c displays the normalized expression levels of representative marker genes of each cluster.

**Fig. 3.**
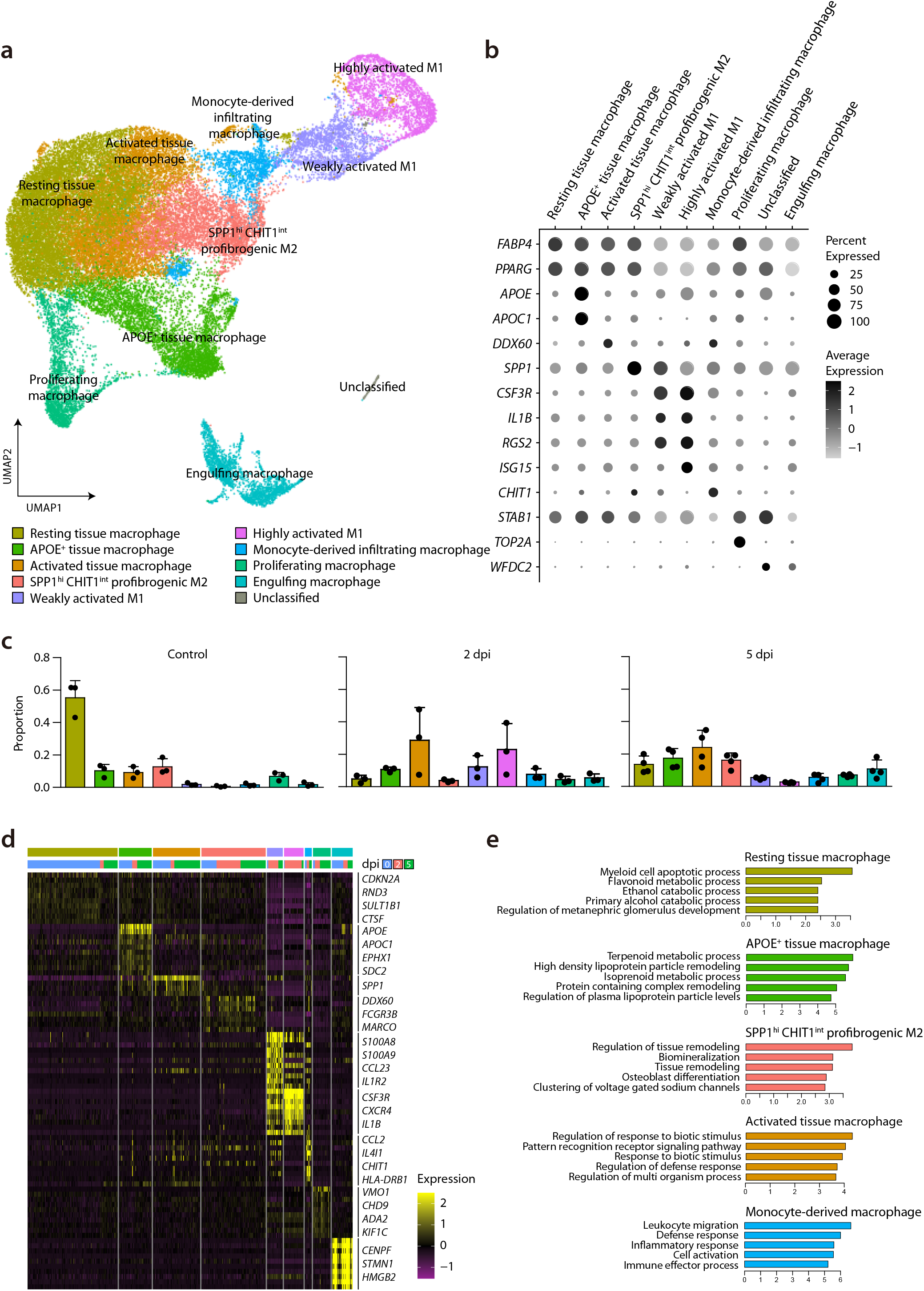
Subpopulation Analysis of Macrophages. a. UMAP plot of the macrophage subpopulations in all groups, colored to show cluster information. b. Ten different clusters and their specific marker gene expression levels, with brightness indicating log-normalized average expression, and circle size indicating the percent expressed. c. Proportion of each macrophage cell type at 0 days post-infection (dpi) (n = 3), 2 dpi (n = 3), and 5 dpi (n = 4). d. Heatmap of cluster-specific differentially expressed genes (DEGs), for each macrophage cell type (n = 9). The color indicates the relative gene expression, and representative genes are shown together. e. Bar plots showing – log10(p value) from enrichment analysis of representative GO biological pathways among resting tissue macrophages, APOE+ tissue macrophages, SPP1hiCHIT1int profibrogenic M2 macrophages, activated tissue macrophages, and monocyte-derived infiltrating macrophages.

The proportion of each lung macrophage subtype underwent distinctive changes. Resting tissue macrophage was the dominant sub-population in control, but was drastically decreased at 2 dpi, and partially recovered at 5 dpi (Fig. 3c and Supplementary Fig. 3b). At 2 dpi, we observed increased proportion of activated tissue macrophages, weakly activated M1 macrophages, highly activated M1 macrophages, and monocyte-derived infiltrating macrophages. At 5 dpi, resting tissue macrophage, APOE^+^ tissue macrophages, activated tissue macrophages, and SPP1^hi^CHIT1^int^ profibrogenic M2 became major populations in proportion, and the proportion of M1 macrophages were lower than 2 dpi. Dynamic changes of the proportions of macrophage subclusters were summarized on UMAP (Supplementary Fig. 3d), and viral-read-containing cells were mainly concentrated in the engulfing macrophage cluster (Supplementary Fig. 3e).

To characterize the subtypes of macrophages in detail, we identified cluster-specific differentially expressed genes (DEGs) (Fig. 3d), and the top 50 DEGs for each cluster were analyzed in terms of gene ontology (GO) biological pathways (Fig. 3e and Supplementary Fig. 3e). DEGs of resting tissue macrophages (the dominant population before SARS-CoV-2 infection) were enriched in GO terms, including “myeloid cell apoptotic process” and metabolism-associated pathways (Fig. 3e). APOE^+^ tissue macrophages had DEGs that were enriched in GO terms mainly associated with lipoprotein metabolism. As expected, DEGs of SPP1^hi^CHIT1^int^ profibrogenic M2 macrophages were prominently enriched in GO terms, including “regulation of tissue remodeling” and biological adhesion, indicating that this subtype is associated with the recovery phase of inflammation. In contrast, activated tissue macrophages and monocyte-derived infiltrating macrophages exhibited DEGs enriched for GO terms associated with activated innate immune response. Supplementary Fig. 3f summarizes the enriched GO terms originated from DEGs of other macrophage sub-clusters. Overall, we defined 10 different subtypes of macrophages in SARS-CoV-2 infection, which displayed extensive heterogeneity.

### Each Macrophage Subpopulation Underwent Transcriptomic Changes Between 2 and 5 Days Post-Infection

Since we observed distinctive proportional changes in the lung macrophage subtypes during SARS-CoV-2 infection (Fig. 3), we next focused on changes in the transcriptome between 2 and 5 dpi in each macrophage subpopulation. Resting and activated tissue macrophages exhibited fewer DEGs than the other macrophage subclusters at 2 and 5 dpi (Fig. 4a and 4b). On the other hand, monocyte-derived infiltrating macrophages showed remarkably increased numbers of DEGs at both 2 and 5 dpi, and exhibited increased expressions of IFN-responsive genes, such as *OAS1*, *ISG15*, and *RSAD2*, at 2 dpi compared to 5 dpi (Fig. 4a). Monocyte-derived infiltrating macrophages exhibited higher expressions of inflammatory markers or mediators, including *HLA-DRB1*, *MRC1*, and *SERPINE2,* at 5 dpi than at 2 dpi. In differentiated macrophage clusters, including M1 and M2 macrophages, the dynamicity of gene expression change was consistently higher at 2 dpi than 5 dpi (Fig. 4b). Weakly and highly activated M1 macrophages showed increased expression of pro-inflammatory genes (including *IL1B*, *CCL8*, and *DUSP1*), while IFN-responsive genes (*OAS1*, *ISG15*, *ISG20*, and *RSAD2*) were upregulated at 2 dpi compared to 5 dpi (Fig. 4b). SPP1^hi^CHIT1^int^ profibrogenic M2 macrophages had different DEGs at 2 dpi, including *SCD*, *CHIT1*, and *IL4I1* (Fig. 4b, right panel). Therefore, monocyte-derived infiltrating macrophages and differentiated M1 and M2 macrophages exhibited increased and distinctive DEG patterns especially at 2 dpi, the peak of viral titer in SARS-CoV-2 infection.

**Fig. 4.**
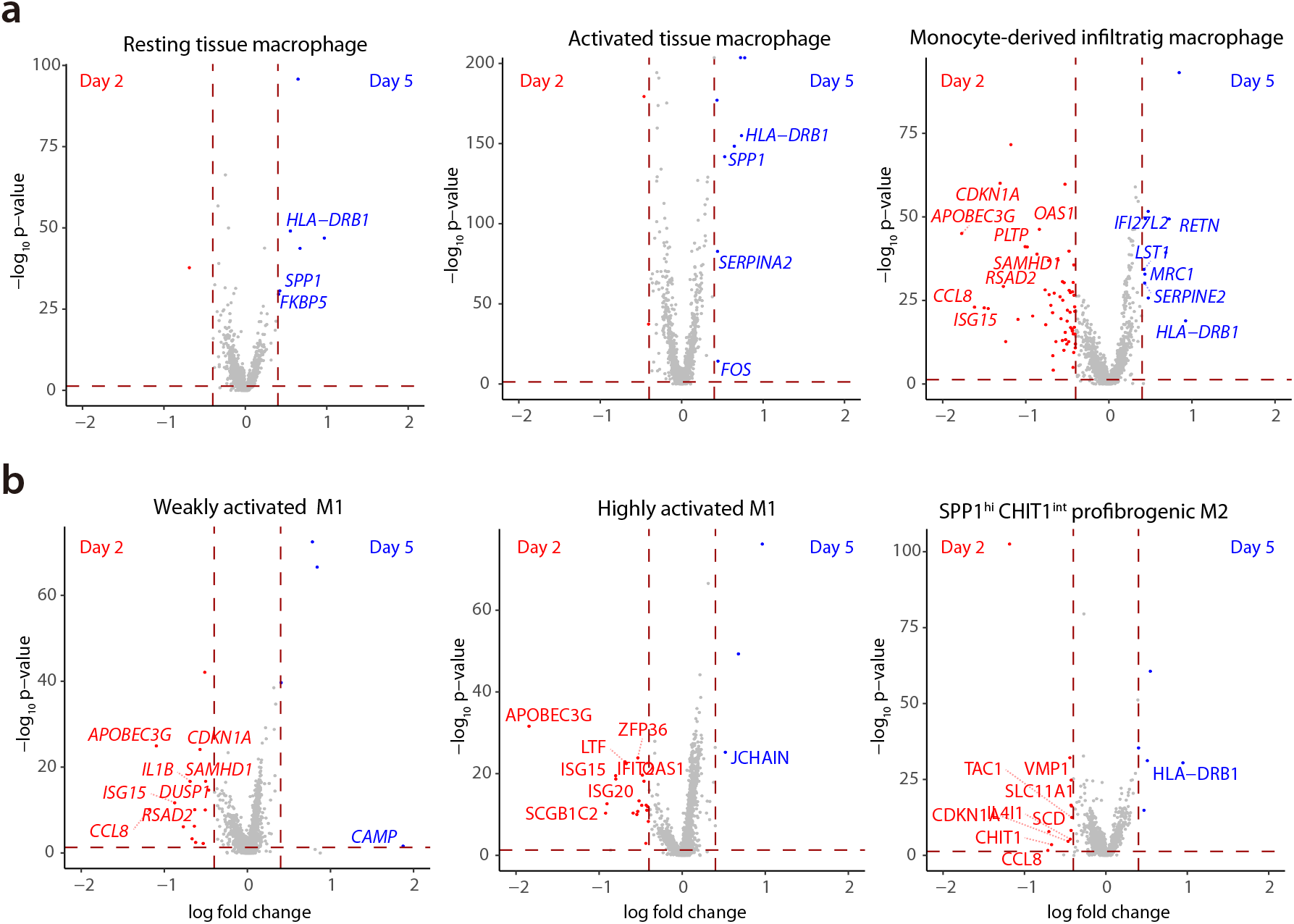
Transcriptomic Changes Between 2 and 5 Days Post-Infection in Macrophage Populations. a, b. Volcano plots showing DEGs between 2 days post-infection (dpi) and 5 dpi among resting tissue macrophages, activated tissue macrophages, monocyte-derived infiltrating macrophages, weakly activated M1 macrophages, highly activated M2 macrophages, and SPP1hiCHIT1int profibrogenic M2 macrophages. Each dot indicates an individual gene. Red indicates a gene that is a significant DEG at 2 dpi, and blue indicates a gene that is a significant DEG at 5 dpi. In the graphs, vertical dashed lines indicate |Log fold change| < 0.4, and horizontal dashed lines indicate p < 0.05.

### RNA Dynamics Revealed Different Spectrums of M1 or M2 Macrophages Originated from Monocyte-derived Infiltrating Macrophages

To further evaluate the RNA dynamics of the macrophage cell subpopulations, we analyzed RNA velocity ^21^. Few kinetics were observed in resting tissue macrophages or activated tissue macrophages, while complex kinetics were formed among monocyte-derived infiltrative macrophages and in both M1 populations (Fig. 5a). To quantify the kinetic dynamics of RNA velocities, we calculated the length of the arrow in Fig. 5a (right panel), which represents the RNA velocities. High velocity levels were formed in both M1 populations. On the other hand, low levels of dynamics were observed in activated tissue macrophages, similar to the resting levels of tissue macrophages, which was consistent with the findings from UMAP embedding (shown in Fig. 5a). Next, we analyzed the direction of the arrow, to investigate the interactions between various clusters. We observed an arrow pointing toward the SPP1^hi^ CHIT1^int^ profibrogenic M2 cluster from the monocyte-derived infiltrating macrophage (Fig. 5a), suggesting that the monocyte-derived infiltrating macrophages significantly contributed to the formation of the SPP1^hi^CHIT1^int^ profibrogenic M2 cluster.

**Fig. 5.**
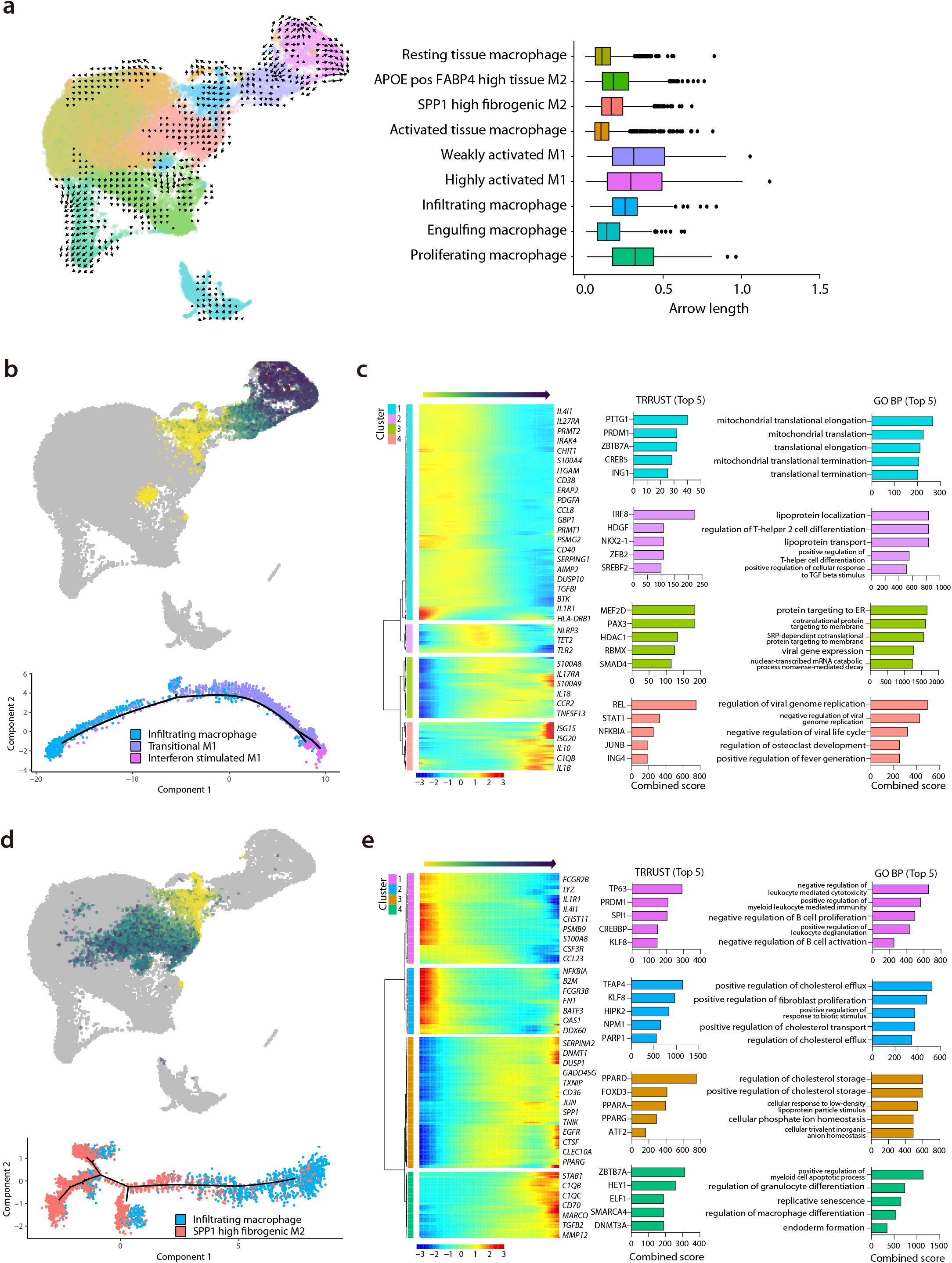
RNA Velocity and Pseudotime Trajectory Analysis from Monocyte-Derived Infiltrating Macrophages to M1 and M2 macrophages. a. Left panel shows UMAP plot of RNA velocity of macrophage subpopulations. Arrow direction and length indicate qualitative and quantitative changes, respectively. Right panel shows box-plots of mean and standard deviation of the arrow lengths in the left panel. b. Pseudotime trajectory initiated from monocyte-derived infiltrating macrophages toward weakly and highly activated M1 macrophages (M1 route). c. Left panel shows relative expression patterns of representative genes in the M1 route plotted along the pseudotime. Color indicates the relative gene expression calculated by Monocle 2. Right panel shows bar plots of the combined scores in the top-five enrichment analysis of the TRRUST database for transcription factor analysis, and representative GO biological pathways in clusters 1–4, as defined in the left panel. d. Pseudotime trajectory initiated from monocyte-derived infiltrating macrophages toward SPP1hiCHIT1int profibrogenic M2 macrophages (M2 route). e. Left panel shows relative expression patterns of representative genes in the M2 route plotted along the pseudotime. Right panel shows bar plots of combined scores in top-five enrichment analysis of the TRRUST database for transcription factor analysis, and the representative GO biological pathways in clusters 1–4, as defined in the left panel.

We next investigated the dynamic transcriptome changes from monocyte-derived infiltrating macrophages to M1 or M2 populations. We found that monocyte-derived infiltrating macrophages were increased during the acute inflammation period, consistent with a previous study^12^. Using pseudotime analysis for single-cell transcriptomics, we traced the dynamic changes of gene expression from infiltrating macrophages to M1 or M2 macrophages^22^. For the trajectory toward M1 macrophages (M1 route) (Fig. 5b; Supplementary Table 5), we defined four distinctive clusters showing modular gene expression changes. We summarized their top 5 associated transcription factors using the TRRUST database^23^, and the top 5 gene ontology biological pathways (GO-BP) (Fig. 5c). Notably, cluster 4 of the M1 route (which was exclusively expressed in highly activated M1 macrophages) showed concurrently increased expressions of *IL1B* and IFN-stimulated genes (*ISG15* and *ISG20*), which were associated with GO terms of enhanced anti-viral activity in the early phase of immune response. These findings indicated that this gene expression change was part of a natural defense mechanism involving M1 macrophage differentiation (Fig. 5b). The highly activated M1 macrophage cluster showed predominant enrichment of pro-inflammatory mediators, including *IL1B* and *CXCL8* (Supplementary Fig. 4a), which was further supported by our results showing that the highly activated M1 was highly enriched with gene sets from severe COVID-19 patients (Supplementary Fig. 4b). These results suggested that the distinct macrophage subpopulation that was potentially derived from monocyte-derived infiltrating macrophages had rapidly undergone SARS-CoV-2-mediated activation of inflammatory macrophage responses.

For the trajectory toward SPP1^hi^CHIT1^int^ fibrogenic M2 macrophages (M2 route) (Fig. 5d; Supplementary table 6), we defined four distinctive clusters and analyzed their features with gene set enrichment analysis, as described in Fig. 5b (Fig. 5e). Cluster 3 of the M2 route showed an increased association with transcription factors of the peroxisome proliferator-activated receptor (PPAR) family (PPAR-δ, PPAR-α, and PPAR-γ) and with pathways associated with cholesterol metabolism. PPAR-γ activation reportedly may drive monocytes toward anti-inflammatory M2 macrophages^24^. Indeed, the next cluster in the pseudotime trajectory, cluster 4 of the M2 route, showed increased expressions of *C1QB*, *C1QC*, *MMP12*, and *TGFB2*, which are known to be key genes of well-differentiated M2 macrophages.

Collectively, the macrophage subpopulations underwent time-dependent and cell-type-specific changes during SARS-CoV-2 infection. These subpopulations exhibited a continuous spectrum of changes, mainly from the monocyte-derived infiltrating macrophages, at the transcriptome level.

### Specific Macrophage Gene Modules from Trajectory Analysis Were Associated with Immune Response to SARS-CoV-2 Infection and Immune-Modulatory Drugs

Next, we compared the dynamically changed macrophage gene modules from M1 and M2 routes with the previously reported transcriptome changes of COVID-19 patients and SARS-CoV-2-infected experimental models^25,26^. Upregulated gene sets determined from postmortem lung tissue of a COVID-19 patient and a SARS-CoV-2-infected mouse were commonly associated with cluster 4 of the M1 route (Fig. 6a). Gene sets from postmortem lung tissue of a COVID-19 patient were also associated with cluster 3 of the M1 route. In contrast, clusters 1 and 2 of the M2 route were highly associated with those three gene sets (Fig. 6b). Therefore, each examined time-point of longitudinal change of macrophage differentiation during SARS-CoV-2 infection encompassed the previous cross-sectional gene sets reported from the COVID-19 patient and experimental model.

**Fig. 6.**
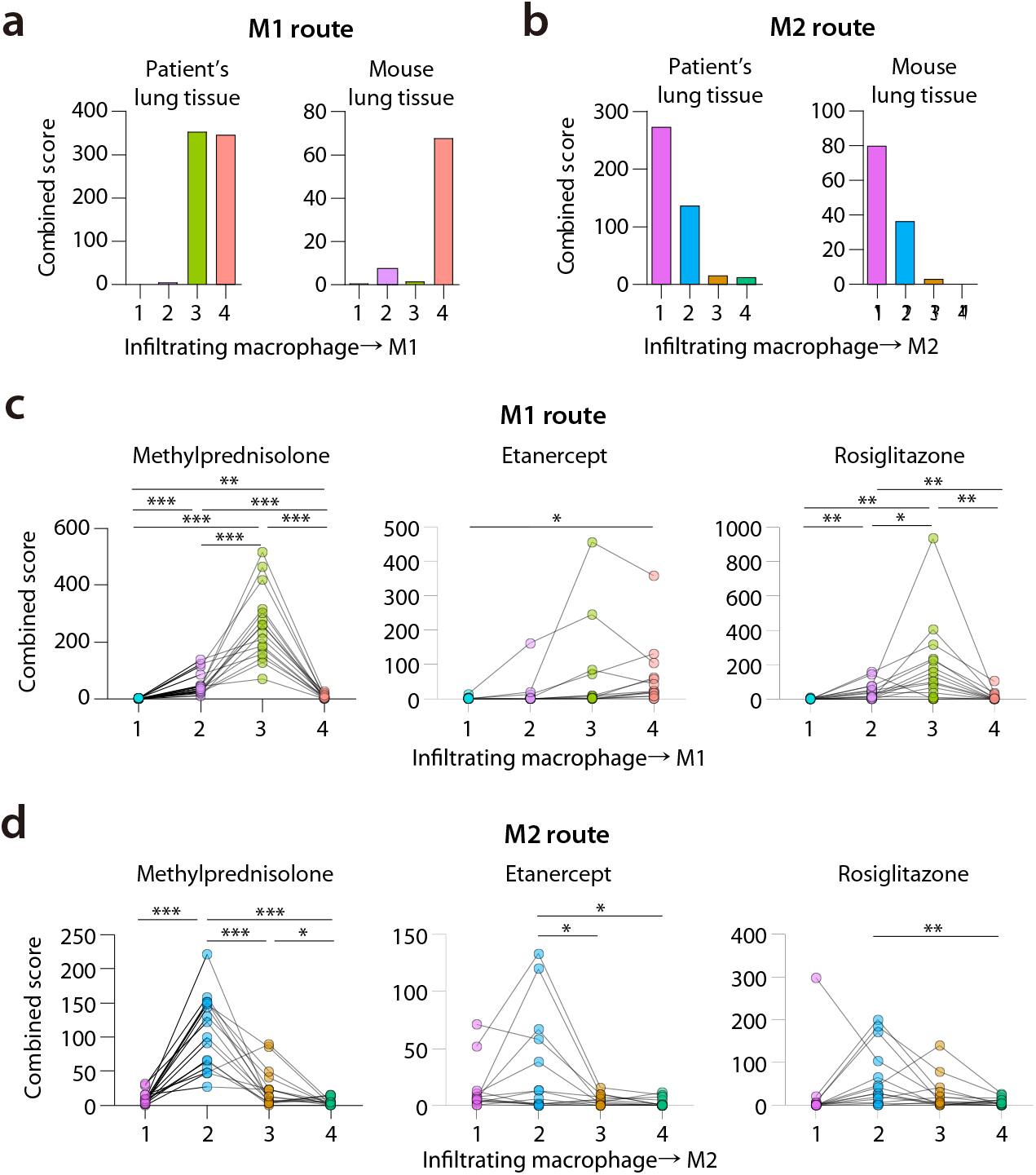
Gene Set Enrichment Analysis of Gene Modules Originated from M1 Route and M2 Route Using Public Datasets Related to SARS-CoV-2 Infection and Immune-Modulatory Drugs. a, b. Gene set enrichment analysis of clusters 1–4 of the M1 route a. and M2 route b. using public transcriptome data, including post-mortem lung tissue from a COVID-19 patient and lung tissue from a SARS-CoV-2-infected mouse. c, d. Gene set enrichment analysis of clusters 1–4 of the M1 route c. and M2 route d. using public transcriptome data, including “Drug Perturbations from GEO down” for methylprednisolone (n = 16), etanercept (n = 14), and rosiglitazone (n = 16). *p < 0.05, **p < 0.01, ***p < 0.001.

Immune-modulatory treatments, including corticosteroids and cytokine-targeted agents, have been considered as a means of regulating hyper-inflammatory responses in COVID-19 patients; however, the exact immunological features of the target cells affected by these treatments is unclear. To evaluate the effect of immune-modulatory drugs on M1 or M2 differentiation, we performed enrichment tests on these trajectory-specific modular gene expressions relative to drug-downregulated gene sets^26^. We found that clusters of the M1 and M2 route were distinctive with regards to transcriptome responses to immune modulatory drugs (Fig. 6c and 6d). For methylprednisolone-induced transcriptome changes, cluster 3 of the M1 route (Fig. 6c, left) and cluster 2 of the M2 route had stronger associations than the other clusters (Fig. 6d, left). For the downregulated gene sets by TNF inhibitor etanercept, it had most predominant association with cluster 4 of the M1 route (Fig. 6c, middle). Etanercept also affected cluster 2 of the M2 route (Fig. 6d, middle). The PPAR-γ agonist rosiglitazone exhibited a pattern of association with clusters of the M1 route (Fig. 6c, right), similar to that of as methylprednisolone, but showed a limited impact on the M2 route, except for cluster 2 (Fig. 6d, right). Our trajectory analysis revealed that most of the transcriptome alterations reported by various sources resembled late clusters of the M1 route and early clusters of the M2 route, and that macrophage-targeting drugs may affect specific stage of M1 or M2 differentiation.

## Discussion

Although recent studies have reported the single-cell transcriptome of BAL fluid cells cross-sectionally obtained from COVID-19 patients, none have used a longitudinal approach along with the natural disease course. In the present study, we investigated single-cell transcriptome changes throughout SARS-CoV2 infection using BAL fluid from a ferret model. We found that specific sub-clusters of NK cells and CD8^+^ T cells exhibited increased responses to IFN, especially at 2 dpi, while their intrinsic cytotoxic properties against viral infection were preserved. More importantly, among macrophages—the major population of BAL fluid cells—we identified 10 different subpopulations that exhibited relative proportion changes from 0 to 5 dpi. The predominant dynamic changes of the transcriptome involved monocyte-derived infiltrating macrophages and differentiated M1/M2 macrophages, especially at 2 dpi. We also observed distinctive and stepwise differentiation from monocyte-derived infiltrating macrophages toward M1 or M2 macrophages.

Our present results included observation of IFN-responsive signatures, regardless of immune cell type, mostly at 2 dpi. The presence of an IFN-responsive signature has also been reported in previous transcriptome studies of SARS-CoV-2 infection^3,4,12^. Data are controversial regarding the relationship between IFN response strength and COVID-19 severity—delayed but robust expression of IFN-associated genes might provoke harmful immunopathology, but their early increase is beneficial ^27^. Our ferret model mimicked SARS-CoV-2 infection with a clinical course of mild severity and spontaneous recovery. Therefore, our findings suggest that prominently increased expression of IFN-responsive genes at 2 dpi might be beneficial in clearing SARS-CoV-2. This observation is further supported by the observed increase of the IFN-stimulated M1 subpopulation.

The BAL fluid cells from our ferret model comprised a diverse subpopulation of macrophages. We annotated 10 different subpopulations among 17 different clusters based on previous single-cell studies of alveolar macrophages^12,24,28–30^. Presence of 0 dpi group provided an interesting contrast with specific features of activated and differentiated macrophages in later phases. The proportion of resting tissue macrophage were near 60% of the macrophage population in control, and drastically decreased at 2 and 5 dpi, suggesting either that this population underwent a change of transcriptomic features towards another population or the infiltration of a new population from circulation. Resting tissue macrophages could have evolved into activated tissue macrophages; however, the increase of activated tissue macrophages was not sufficient to fully explain the decreased proportion of resting tissue macrophage. Notably, the increased RNA velocity of infiltrating and M1/M2 macrophages indicated that these were the major populations that underwent dynamic changes after SARS-CoV-2 infection. Here, we found that with regards to the changing macrophage populations, resting tissue macrophages decreased after inoculation but were not restored later, and M2 macrophages were increased and remained a major population from 2 to 5 dpi. These findings indicate that during the viral resolution phase, an active repair process is underway rather than complete recovery to pre-infection status.

Immuno-modulatory treatments—including corticosteroids and targeted agents, such as Janus kinase inhibitors—have been considered to regulate hyper-inflammatory responses in COVID-19 patients^9,10,27^. However, to apply such treatments in heterogeneous COVID-19 patients, we must understand the exact features and proportions of the target immune cell populations that will be affected. Along the transcriptome continuum of monocyte-derived infiltrating macrophages to M1 macrophages (the M1 route), we found that the later clusters, similar to highly activated M1 macrophages, were enriched in gene sets related to treatment with corticosteroid, TNF inhibitor, and PPAR-γ agonist. Additionally, along the M2 route, the earlier phase rather than the later phase of the transcriptome features was enriched in gene sets from a COVID-19 patient’s lung tissue, other model systems, and in medication-downregulated gene expression changes. These findings suggest that those medications may contribute to proper suppression of the M1-associated hyper-inflammation response without significantly affecting the M2-associated resolution process. Corticosteroid therapy reduces mortality in cases of severe pneumonia^31^, and the beneficial role of dexamethasone in hospitalized COVID-19 patients has also been reported recently^32^. Our current findings support the potential benefits of proper immune suppression, and elucidate the exact subpopulations affected by these macrophage-affecting medications.

Overall, our present study provides fundamental information regarding the immune response dynamics provoked by SARS-CoV-2 infection, as well as a detailed description of the natural course and changes of macrophages in the ferret model.

## Methods

### Experimental Animals

Experiments were performed using 14-to 20-month-old female ferrets (n = 10, ID Bio Corporation, Cheongju, Korea) that were serologically negative for influenza A viruses (H1N1 and H2N2), MERS-CoV, and SARS-CoV. Ferrets were maintained in the isolator (Woori IB Corporation, Daejeon, Korea) in BSL3 of Chungbuk National University. All ferrets were group housed with a 12-h light/dark cycle, and allowed access to food and water. After two days of adaption to BSL3 conditions, the ferrets were intranasally inoculated with phosphate-buffered saline (PBS) (n = 3) or 10^5.8^ TCID_50_/mL of NMC-nCoV02 (n = 7), while under anesthesia with ketamine (20 mg/kg) and xylazine (1.0 mg/kg). All animal studies were conducted following protocols approved by the Institutional Animal Care and Use Committee (IACUC) of Chungbuk National University (Approval number CBNUA-1352-20-02).

### Virus and Cells

SARS-CoV-2 strain NMC-nCoV02 (reference, Cell host & Microbe) was propagated in Vero cells in Dulbecco’s Modified Eagle Medium (DMEM; Gibco, Grand Island, NY) supplemented with 1% penicillin/streptomycin (GIBCO) and TPCK-treated trypsin (0.5 μg/mL; Worthington Biochemical, Lakewood, NJ) in a 37°C incubator with 5% CO_2_ for 72 h. The propagated virus was then stored at −80°C, and used as the working stock for animal studies. The 50% tissue culture infective dose (TCID_50_) was determined via fixation and crystal violet staining.

### Harvesting Bronchoalveolar Lavage Cells

At 2 and 5 dpi, respectively, three and four ferrets were euthanized, and bronchoalveolar lavage fluid (BALF) was collected. As a control group, the three PBS-treated ferrets were euthanized at 2 dpi and BALF was collected. Briefly, with the ferret positioned in dorsal recumbency, 30 mL of cold sterile PBS solution containing 5% fetal bovine serum (FBS) was injected through the tracheal route and then collected. This collected lavage fluid was centrifuged at 400 × g for 10 min at 4°C. Then the supernatant was removed, and the cell pellet was suspended in 5 mL 10X RBC lysis buffer (Thermofisher, cat. no. 00-4300-54) diluted 1:10 with distilled water, followed by a 10-min incubation at room temperature. After the RBC lysis reaction, 20 mL of 1X PBS was added to stop the lysis reaction, followed immediately by centrifugation at 500 × g for 5 minutes at 4°C. Then the supernatant was removed, followed by cell number and viability analyses.

### Virus Isolation From the Lungs of Infected Ferrets

The virus titers in collected lung tissues were determined by TCID_50_ in Vero cells. Briefly, lung tissue samples were homogenized in an equal volume (1 g/mL) of cold 1X PBS containing 1% penicillin/streptomycin (GIBCO). Tissue homogenates were centrifuged at 3000 rpm for 15 min at 4°C, and then the supernatants were serially diluted (10^−1^ to 10^−8^) in DMEM. Dilutions of each sample were added to Vero cells, followed by a 2-hour incubation. Next, the media (DMEM) was changed, and the cytopathic effects (CPEs) were monitored for 4 days. We determined the TCID_50_ through fixation and crystal violet staining.

### Histology

Lung tissue samples were collected at 2 and 5 dpi, incubated in 10% neutral-buffered formalin for fixation, and then embedded in paraffin following standard procedures. The embedded tissues were sectioned and dried for 3 days at room temperature. Then the tissue sections were placed on glass slides, stained with hematoxylin and eosin (H&E), and compared with PBS control group. Slides were viewed using an Olympus IX 71 (Olympus, Tokyo, Japan) microscope, and images were captured using DP controller software.

### scRNA-Seq Analysis: Basic Quality Control

Reference sequence and gene information were downloaded from the Ensembl database (MusPutFur1.0, under accession number GCF_000215625.1), and then annotated with human ortholog genes using the same database (Biomart database, GRCh38). The SARS-CoV-2 sequence was downloaded from NCBI GenBank (Wuhan-Hu-1, a widely used reference sequence, under accession number NC_045512). Reference genome information was pre-processed for single-cell data processing using mkref (Cell Ranger 10x genomics, v3.0.2), and the fastq files were generated through the process of demultiplexing the sequenced data (Cell Ranger). Next, the reads were aligned to the ferret–virus combined reference genome, and the aligned read data were analyzed using Seurat R package v3.1.5 ^33^. Based on the characteristics of inflammatory tissue and the assumption that viral transcripts can present in dying cells, we did not exclude low-quality cells from the analysis. Ambient RNAs were examined and adjusted using SoupX (https://doi.org/10.1101/303727), and were present in 1– 3% of each sample, indicating that the samples were relatively clean/washed. We also excluded doublets perceived based on dual expression of cell-type specific gene expression markers, which were dominant in the cluster “Doublet.” Despite high variability in the number of UMIs detected per cell, most cells were enriched with UMIs within a reasonable range (interquartile range: 2,455 to 12,764).

In each cell, gene expression was normalized and scaled using the SCTransform algorithm ^34^. Dimensional reduction and visualization were performed via principal components analysis (PCA) and Uniform Manifold Approximation and Projection (UMAP)—using the top 20 principal components (PCs) for whole cell types, 5 PCs for NK and CD8 T cells, and 13 PCs for monocyte/macrophage cell types—with parameters of min.dist = 0.2, and n.neighbor = 20. Lastly, the cells were clustered by unsupervised clustering, using the default pipeline of the Seurat package (resolution = 0.4 for whole cell types, 0.3 for NK cells, 0.2 for CD8 T lymphocytes, and 0.6 for monocytes/macrophages). We observed two polymorphic genes that significantly affected the clustering of a subset of macrophages by samples: *HLA-DQA1* and ENSMPUG00000007244, the latter of which is putative *HLA-DQB1* or *HLA-DQB2*, and has a DNA sequence that overlaps 78.03–78.81% with human *HLA-DQB1* or *HLA-DQB2*. We removed these two genes from the count matrix and re-processed, and found that the batch effect was resolved.

### Marker Detection and Differential Expression Analysis

To identify marker genes, we selected genes in each cluster that were upregulated relative to the other clusters, based on the Wilcoxon rank-sum test in Seurat’s implementation (FindAllMarkers function), with a >0.25 log fold change compared with the other clusters and a Bonferroni-adjusted p value of <0.05. To investigate the dynamic changes in gene expression in certain cell clusters, we tested differentially expressed genes, using the Wilcoxon rank-sum test (Fig. 4a and 4b). Gene names that had a human ortholog were marked when the p value was <0.05, and the absolute value of the log2 fold change was >0.4.

### GO and Pathway Enrichment Analyses

As shown in Fig. 3e and 3f, cluster-specific expression markers were subjected to Gene Ontology (GO) enrichment analysis ^35^, which is based on the performance of Fisher’s exact test on curated gene sets annotated according to the gene ontology consortium in the biological process category. Ontology terms associated with T cells and eosinophils, and near-duplicated terms, were removed using a custom script, with the following exclusion criteria: GO terms, including ‘T_HELPER|T_CELL’, ‘EOSINOPHIL’, ‘POSITIVE’ or ‘NEGATIVE’. For each cluster, the top 50 genes (prioritized by fold change when comparing each cluster with the rest) were subjected to the enrichment test. Genes that were expressed in >80% of cells in the rest of the clusters were excluded.

To predict transcription factors that might drive macrophage differentiation in pathology, the same enrichment test was performed using the TRRUST transcription factor-target gene database ^23^. To identify potential drugs for controlling macrophage differentiation, the same test was performed using a manually curated dataset based on ‘Drug Perturbations from GEO down’ in enrichR ^26^. This dataset originated from the transcriptome of samples treated with methylprednisolone (GSE490), etanercept (GSE11903, GSE36177, GSE41663, GSE47751, and GSE7524), and rosiglitazone (GSE11343, GSE1458, GSE36875, GSE7193, GSE5509, GSE5679, GSE7035, GSE10192, GSE2431, GSE21329, and GSE35011).

### RNA Velocity

To investigate the characteristics of RNA dynamics among macrophages in the ferret model, we analyzed RNA velocity based on modeling gene expression induction and repression using spliced and unspliced reads. This technique was previously demonstrated to be feasible in a 3 captured single-cell RNA sequencing library using the velocyto tool ^21^. Spliced and unspliced reads were counted using the run10x command in the velocyto tool with default options. The count matrixes were filtered using velocyto’s standard pipeline, with min.max.cluster.average parameters of 0.08 for the spliced read count matrix, and 0.06 for the unspliced read count matrix. Among a macrophage/monocyte population of 40,241, 5,000 cells were randomly selected, with pooling of the 20 nearest neighbors in the spliced/unspliced count matrix. Through this process, the cell distance matrix was derived from Seurat’s shared neighborhood network matrix with default parameters (FindNeighbors function). Velocity estimation was conducted using the options of deltaT = 1, fit.quantile = 0.05, and kCells = 1 (as k-nearest neighbor pooling was already performed before the random sampling of 5000 cells).

### Analysis of Dynamic Transcriptome Changes Accompanying M1 and M2 Differentiation

To investigate the dynamic changes along the M1 and M2 differentiation pathway, we exported related cell clusters for monocle’s standard analysis process. The related clusters included weakly activated M1, highly activated M1, and monocyte-derived infiltrating macrophages for the M1 pathway; and monocyte-derived infiltrating macrophages and SPP1h CHIT1int profibrogenic M2 for the M2 pathway. Briefly, CellDataSet objects were built based on normalized count (SCTransform), and then processed using estimateSizeFactor and estimateDispersions function (default option), detectGenes (with the min_expr = 0.1 option), setOrderingFilter and reduceDimension (with options of max_components = 3, and method = “DDRTree”), orderCells (default option), and plot_cell_trajectory (default option). Trajectory-specific genes were grouped into four clusters using hierarchical clustering. Finally, each cluster was subjected to further enrichment analysis for transcription regulation or ontology-based analysis.

## Statistical Analysis

The statistical significance of the combined scores from GSEA results were assessed by paired T test. Data plotting, interpolation and statistical analysis were performed using GraphPad Prism 8.2 (GraphPad Software, La Jolla, CA). Statistical details of experiments are described in the Figure legends. A p value less than 0.05 is considered statistically significant.

## Author Contributions

Conceptualization, JSL, KCY, and SHP; Methodology, JSL, JYK, KY, and YIK; Investigation, JSL, JYK, KY, YIK, SJP, SHP, YSJ, YKC, and SHP; Resources, YKC, YSJ, and SHP; Writing, JSL, JYK, KY, YKC, and SHP; Review & Editing, JSL, JYK, KY, SHP, YKC, and SHP; Supervision, JSL, KCY, and SHP; Funding, YKC, and SHP.

## Author Information

The authors declare no competing financial interests: details are available in the online version of the paper. Readers are welcome to comment on the online version of the paper. Correspondence and requests for materials should be addressed to J.S.L., Y.K.C., S.H.P. All raw and processed data have been deposited in the GEO database under accession number GSEOOOOOO.

## Acknowledgements

We thank Eui-Cheol Shin (KAIST) and Jongeun Park (KAIST) for valuable discussion and productive comments. This work was supported partially by the grant from the Korea Health Technology R&D Project through KHIDI funded by the Ministry of Health & Welfare (HI20C0546 to S.H.P), the National Research Foundation of Korea (NRF) grant (NRF-2019R1A2C2005176 to S.H.P), and by Mobile Clinic Module Project funded by KAIST (to S.H.P). This work was also supported by NRF grants (NRF-2020R1A2C 3008339 to Y.K.C) and for Research Center for Severe Pulmonary Disease (2020R1A5A2017476 to Y.K.C).

## Code Availability

For all data analyses, we used publicly available software.

## Supplementary Materials

**Supplementary Figure 1.**
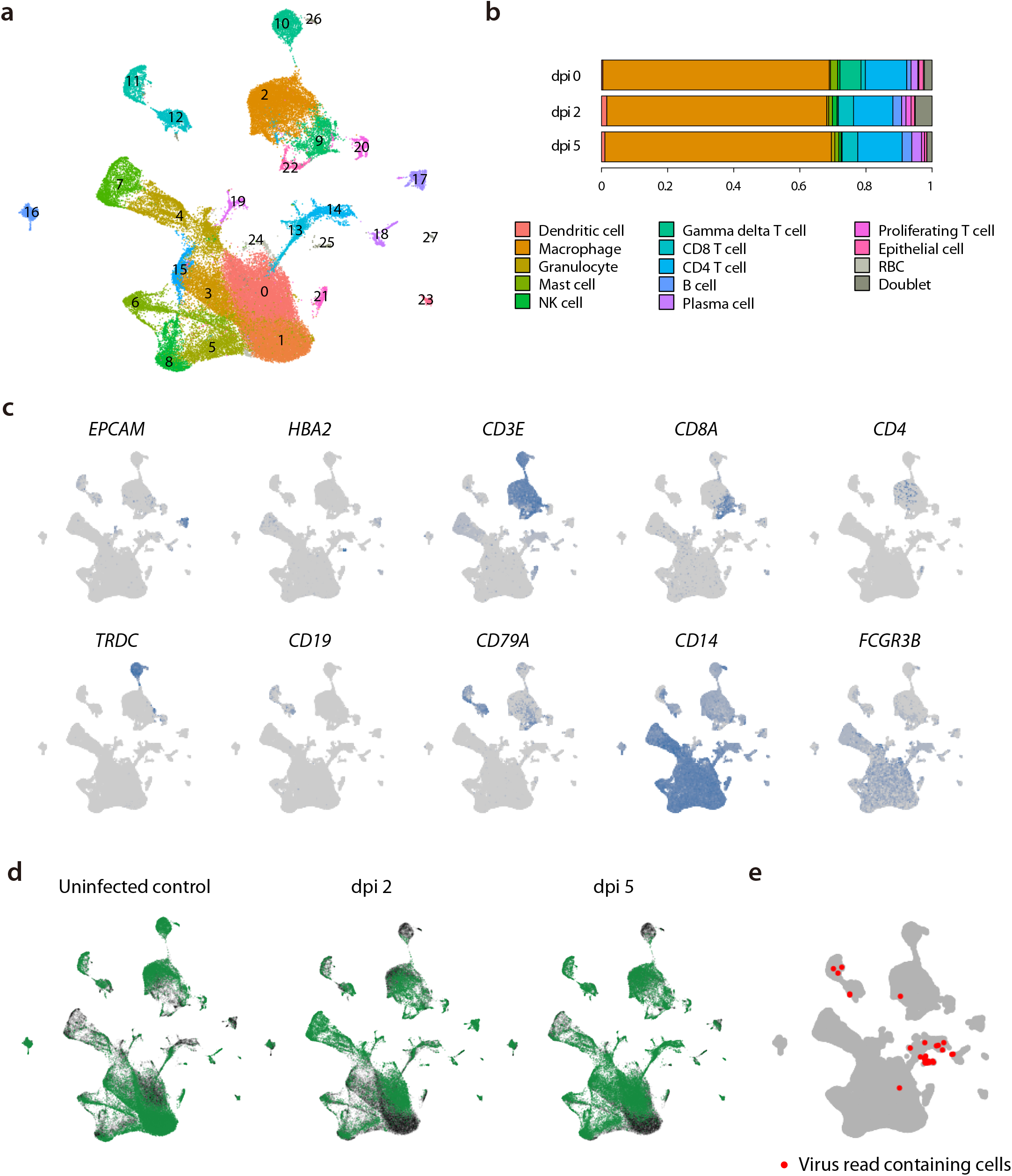
(related to Fig. 1) a. UMAP plot colored according to cluster. b. Proportion of each cell type at 0 days post-infection (dpi) (n = 3), 2 dpi (n = 3), and 5 dpi (n = 4). c. UMAP plots showing normalized expression of known markers. d. UMAP plot, with color density reflecting the distribution of cells at 0, 2, and 5 dpi. e. UMAP plot of virus-read-containing cells (red dots).

**Supplementary Figure 2.**
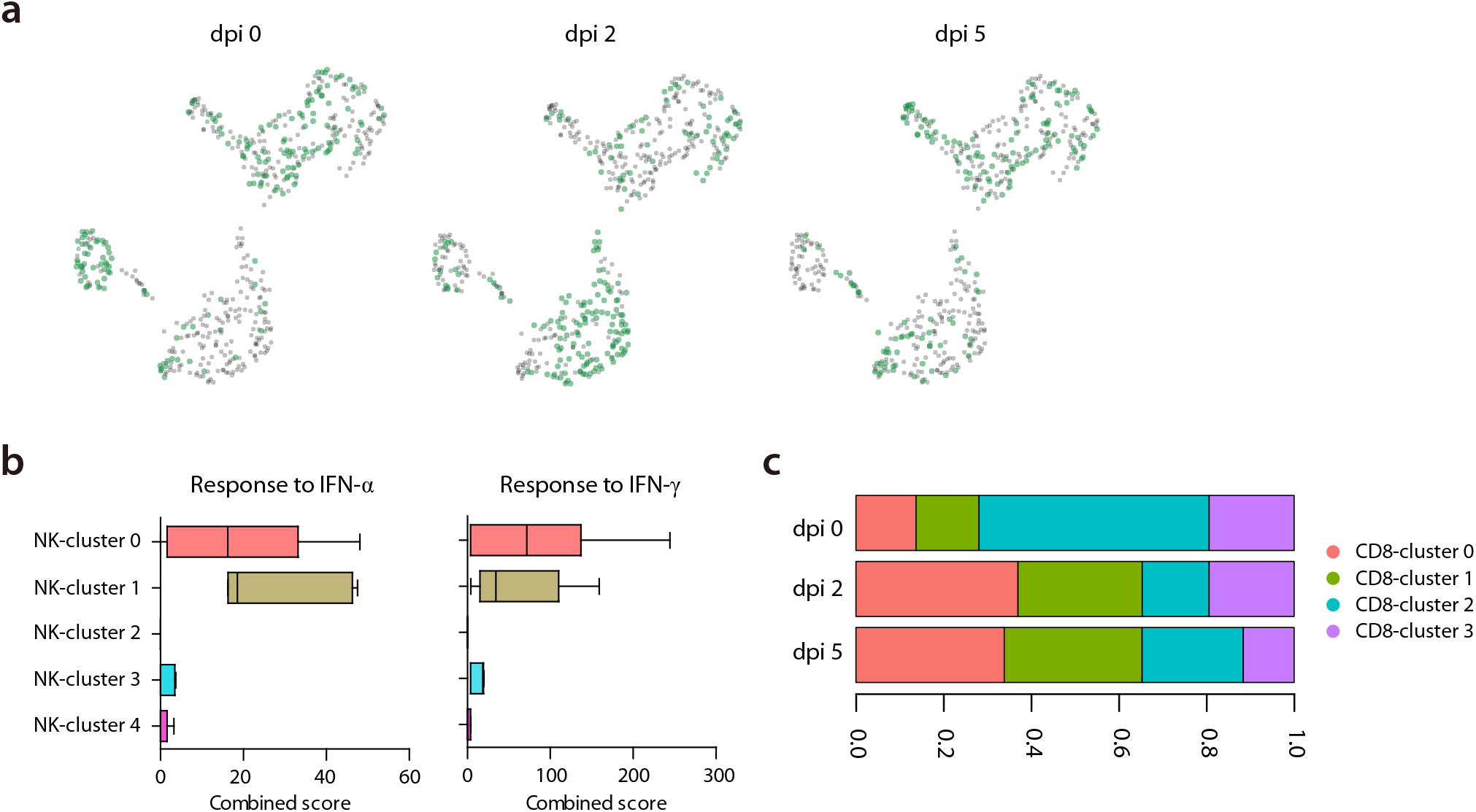
(related to Fig. 2) a. UMAP plot, with color density reflecting the distribution of NK cells at 0, 2, and 5 days post-infection (dpi). b. Box-plots showing the results of gene set enrichment analysis on five NK cell clusters using two gene sets: “Response to IFN-α” (left) and “Response to IFN-γ” (right). c. Proportion of each cell type in CD8+ T cell clusters at 0 dpi (n = 3), 2 dpi (n = 3), and 5 dpi (n = 4).

**Supplementary Figure 3.**
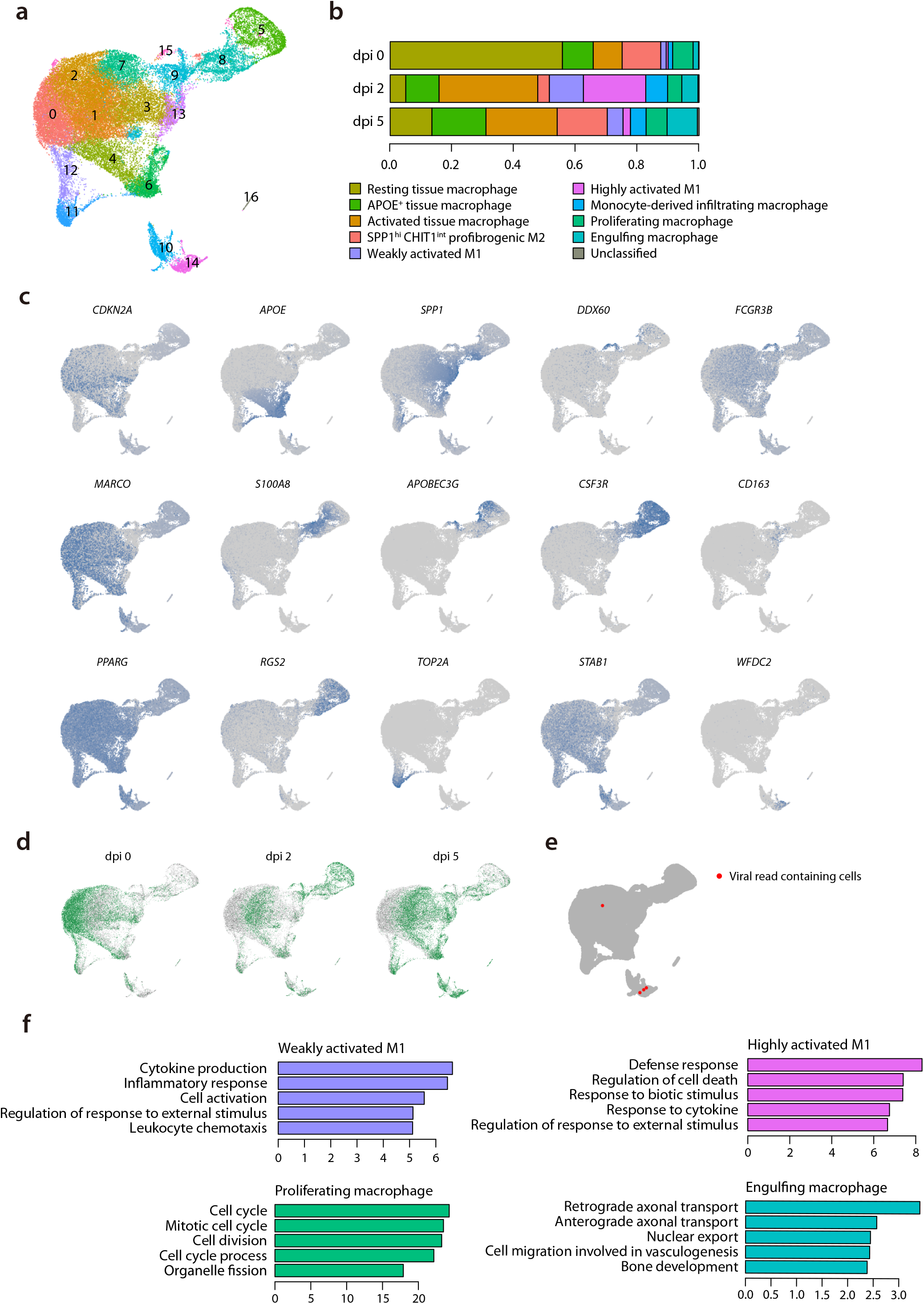
(related to Fig. 3) a. UMAP plot of macrophage subpopulations, colored according to clusters. b. Proportion of each macrophage subpopulation at 0 days post-infection (dpi) (n = 3), 2 dpi (n = 3), and 5 dpi (n = 4). c. UMAP plots showing normalized expression of known markers of macrophage subpopulations. d. UMAP plot with color density reflecting distribution of macrophage subpopulations at 0, 2, and 5 dpi. e. Virus read containing cells (red dot) in UMAP plot of macrophage subpopulations. f. Bar plots showing –log10(p value) in top-five enrichment analysis of representative GO biological pathways among weakly activated M1 macrophages, highly activated M1 macrophages, proliferating macrophages, and engulfing macrophages.

**Supplementary Figure 4.**
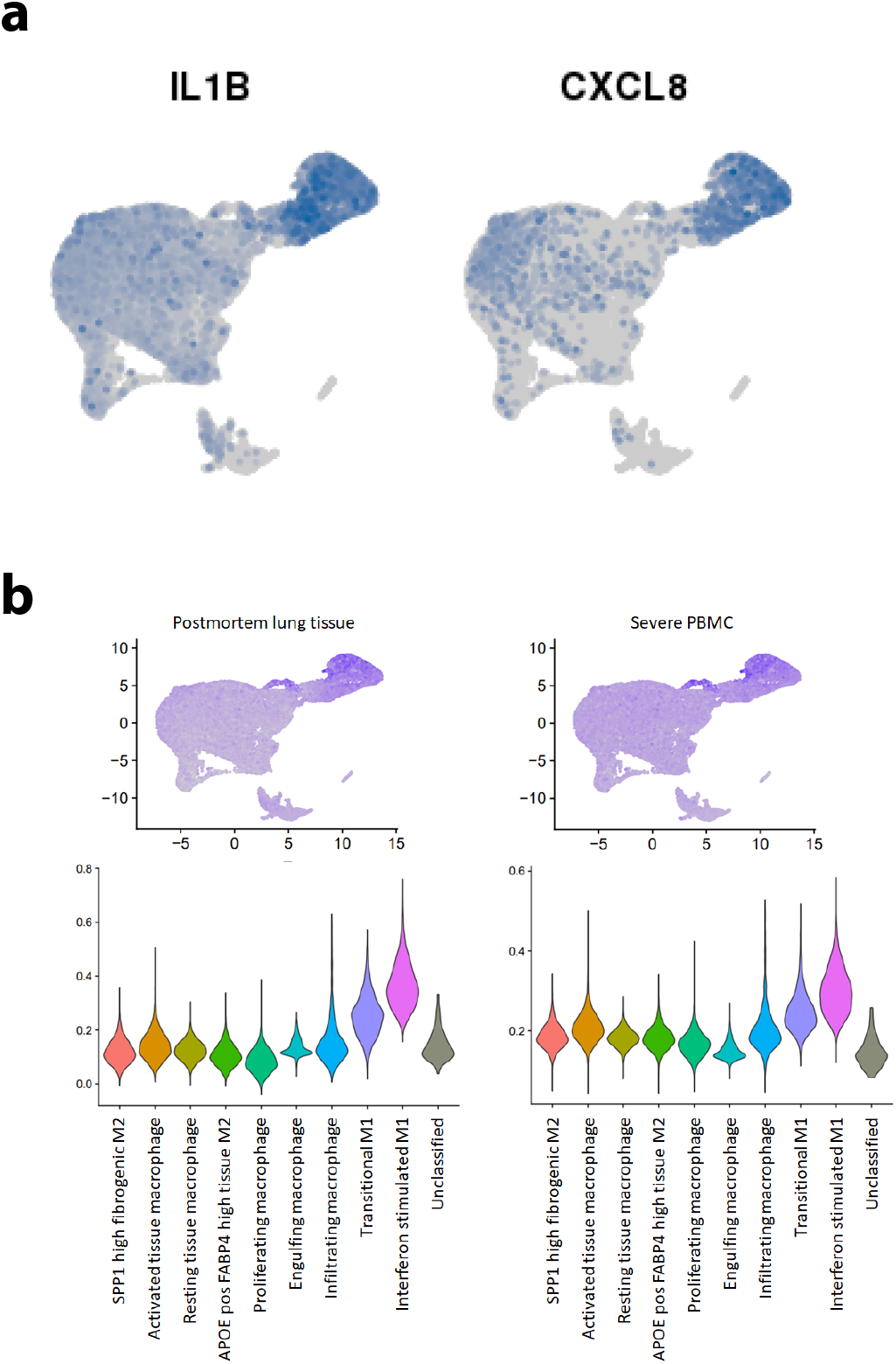
(related to Fig. 5) UMAP plots showing normalized expressions of IL1B and CXCL8 in the macrophage subpopulations.

**Supplementary Table 1**. List of Marker Genes for Each Cluster of Total BAL Fluid Cells

**Supplementary Table 2**. List of Marker Genes for Each Subcluster of NK Cells

**Supplementary Table 3**. List of Marker Genes for Each Subcluster of CD8+ T Cells

**Supplementary Table 4**. List of Marker Genes for Each Subcluster of Macrophages

**Supplementary Table 5**. List of Genes Upregulated in Clusters 1–4 of M1 Route Pseudotime

**Supplementary Table 6**. List of Genes Upregulated in Clusters 1–4 of M2 Route Pseudotime

